# Interplay of CaHVA and Ca^2+^-activated K^+^ channels affecting firing rate in perineuronal net disruption

**DOI:** 10.1101/2023.06.05.543694

**Authors:** Kine Ødegård Hanssen, Torbjørn Vefferstad Ness, Geir Halnes

**Affiliations:** Department of Physics, University of Oslo, Oslo, Norway; Centre for Integrative Neuroplasticity, University of Oslo, Oslo, Norway; Department of Physics, Norwegian University of Life Sciences, Ås, Norway

**Keywords:** Perineuronal nets, CaHVA channel, SK channel, BK channel, Firing rate, PV cells, Fast-spiking interneurons, One-compartment models of neurons

## Abstract

Perineuronal nets (PNNs) are extracellular matrix structures consisting of proteoglycans crosslinked to hyaluronan. They wrap around subgroups of individual neurons in the brain, primarily parvalbumin positive inhibitory neurons. The nets have been found to affect conductances and activation curves of certain ion channels, including Ca^2+^-activated K^+^ and high-voltage-activated Ca^2+^ channels. We studied how PNN related parameters affected the firing rate of one-compartment neuron models. We found that the direct effect of the CaHVA current on firing rate was small, while it had a much larger indirect effect on firing rate through initiation of the SK current. Upregulation of the SK conductance similarly had a pronounced effect on the firing rate. The SK currents therefore acted as the main determinant of firing rate out of these two mechanisms. We shifted the CaHVA channel activation by 14.5 mV and increased the SK conductance by a factor of 3.337, consistent with experimental findings on PNN breakdown in the literature. We studied this in nine different models and found a reduction in firing rate in some, but not all, of these models.

## Introduction

Perineuronal nets (PNNs) are structures made up primarily of glycosaminoglycans and crosslinkers. The nets enwrap neurons of various types in various parts of the brain, but tend to be located on parvalbumin positive (PV) neurons [31]. PNNs have been linked to memory and mature at the end of a period of increased plasticity [30, 29]. They have also been indicated to protect their ensheathed neurons from neurotoxicity [31] and pose a physical barrier to ion transport [31, 16]. Furthermore, PNN digestion has been shown to reduce the firing rate of PV neurons [28, 1, 22, 5].

Tewari et al. [28] found an increased capacitance *c*_m_ and a reduced firing rate *f* in PV neurons when PNNs were disrupted. However, trying to replicate their experiments [28] using biophysically detailed models, we found that the observed change in *c*_m_ alone could not account for the experimentally observed change in firing rate [17]. Other studies have indicated that that PNNs affect ionic currents [6, 32, 34, 10], and ionic concentrations [24, 3], which should affect the reversal potentials. In a previous work, we managed to reproduce the reduction in *f* observed by Tewari et al. [28] by combining the reported increase in *c*_m_ with changes in conductances and reversal potentials [17].

The literature suggest that both the activation of high-voltage-activated Ca^2+^ (CaHVA) current [32, 20] and the small-conductance Ca^2+^-activated K^+^ (SK) current [6] are affected by the presence of PNNs or its components. In our previous study [17], we found that the SK current caused a reduction in *f* and could help explain the reduction in *f* observed experimentally [28]. Since SK channel activation is triggered by Ca^2+^, CaHVA will affect firing both directly through influx of Ca^2+^ and indirectly through activating the SK channel. The effect of this interplay of CaHVA and SK on *f* following PNN degradation warrants further study.

Vigetti et al. [32] found that the activation of high-voltage-activated Ca^2+^ (CaHVA) current in xenopus photoreceptors was greatly affected by the presence of chondroitin sulphates (CS), a key component of the PNNs. The presence of CS in solution shifted the activation curve towards lower voltages. Removal of the nets should therefore shift the activation curve to higher voltages. Possibly related to this, Kochlamazashvili et al. [20] found that the presence of hyaluronan (HA), another net component, increased the amplitude of Ca^2+^ current. While this could be an effect of an increased maximal Ca^2+^ conductance, it may also be due to the nets shifting the activation curves to lower voltages as observed by Vigetti et al. [32], causing the channels to open at lower voltages.

Dembitskaya et al. [6] found that removal of PNNs with chondroitinase ABC (chABC) yielded an increase in the small-conductance Ca^2+^-activated K^+^ (SK) current, which is known to be afterhyperpolarizing [9]. They found that new dendritic spines appeared when PNNs were removed and no longer blocked the growth of these spines. They hypothesized that the new dendritic spines up-regulate 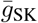by increasing the SK-channel expressing membrane area [6].

Due to the low intracellular Ca^2+^-concentration, Ca^2+^ currents tend to be inward and hence depolarize the membrane. The K^+^ equilibrium potential is low, so K^+^ tends to flow out of the cell and repolarize or hyperpolarize the membrane potential. Hence, the presence of a Ca^2+^ channel might increase the firing as Ca^2+^ acts to bring the membrane potential closer to the firing threshold, while K^+^ channels such as SK have the opposite effect. When both mechanisms are present, these two mechanisms may interact in ways that might be sensitive to the relative values of the conductances. Therefore, we have looked at the effect of the CaHVA and SK channels on firing.

We have two research objectives. The first objective is to study how alterations to CaHVA and SK channels in PV cells affect the firing rate *f*. To address this, we first study how changing the maximal conductances 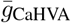 and 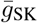 affect *f* in a one-compartment model with CaHVA and SK channels taken from a PV neuron model in the Allen Institute’s database [12]. We refer to this as our main model. Then, we study the effect of introducing a shift in the CaHVA channel activation on *f* in this model. We find that increasing 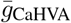 has limited effect on *f* unless the SK conductance is non-zero, while increasing 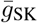 has a pronounced effect on *f*. Shifting the CaHVA channel activation also has limited effect on *f* unless 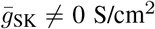 These findings strongly suggest that effects on *f* are mainly due to *I*_SK_.

The second research objective is to model PNN disruption according to literature and examine whether the resultant change in *f* is consistent with observations of *f* on PV cells with removed nets. To achieve this goal, the CaHVA activation is shifted by 14.5 mV [32] and 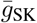 is increased by a factor of 3.337 [6]. We find that the changes reported in the literature induce an increase in *f*, so that altering the CaHVA and SK currents together cannot explain the reduced firing associated with PNN breakdown in our main model. To check if this finding generalizes to other models, we construct eight additional one-compartment models having different combinations of CaHVA and Ca^2+^-activated K^+^ channels, which we adopt from other PV neuron models. We find that parameter changes associated with PNN breakdown increase *f* in some of the models, and reduce *f* in others, something that again highlights the complexity in the interplay between CaHVA and Ca^2+^-activated K^+^. Although our main model suggests otherwise, we thus cannot rule out the possibility that changes in CaHVA and SK channels can explain the observed reduction in *f* associated with PNN degradation.

## Methods

In order to get firing characteristic of PV cells, we use a one-compartment base model that only contains a Na^+^ current *I*_naf_ and a K^+^ current *I*_kaf_ from Hjorth et al. [18], along with a general leak current *I*_L_. The action potentials (APs) that this model produce are narrow, which is characteristic of PV cells [2].

The main model used in this study is built upon the base model, but with additional currents *I*_CaHVA_ and *I*_SK_ that were taken from a PV cell model by the Allen Institute for Brain Science [12]. More specifically, we used conductance values and mechanisms from the perisomatic model of the neuron with CellID 471077857 in the Allen Brain Atlas database, lowering the maximal CaHVA conductance by a factor of five in order to obtain the high firing frequency characteristic of the PV cells [28, 22]. The currents in the main model are on the form

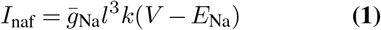

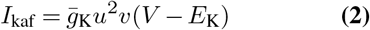

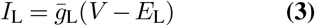

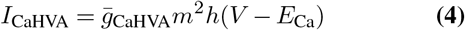

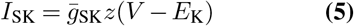

The variables of activation and inactivation all vary according to

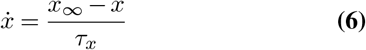

where *x* ∈ {*x l, k, u, v, m, h, z*} is the activation or inactivation variable *x*_*∞*_ and 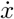 is its time derivative. *x* and *τ* are variables that depend on either *V* or [Ca^2+^]_in_.

As the CaHVA and SK channels are the main focus of this work, the equations for their kinetics are listed below. The expression for the corresponding variables for *I*_naf_ and *I*_kaf_ can be found in Hjorth et al. [18].

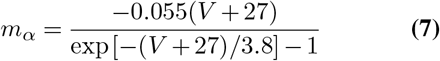

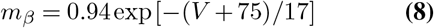

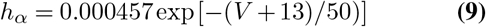

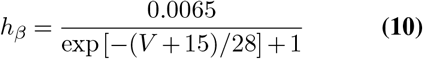

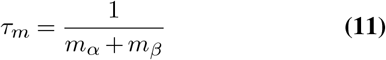

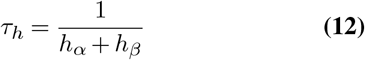

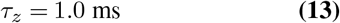

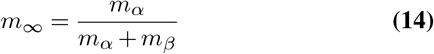

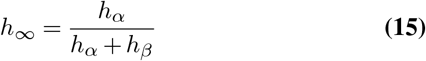

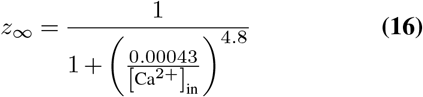

The Ca^2+^ reversal potential *E*_*ca*_ depends strongly on the intra- and extracellular Ca^2+^ concentrations as these vary a lot [27]. In order to obtain reliable results, the Ca^2+^ concentrations varied during the simulations to vary according to

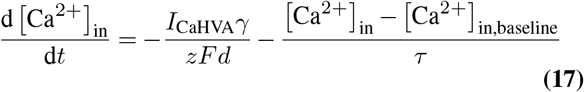

where *γ* is the percent of free Ca^2+^ and *τ* is the rate of Ca^2+^ decay. *z* = 2 is the valence of the Ca^2+^ ion, *F* is Faraday’s constant and *d* = 0.1 *μ*m is the depth of an imaginary submembrane shell [7]. [Ca^2+^]_in,baseline_ is the baseline intracellular Ca^2+^ concentration. *E*_Ca_ was calculated by NEURON at each time step using these concentrations [12].

In order to yield appropriate Ca^2+^ dynamics, the parameters in Eq. 17 were set to *γ* = 0.2 and *τ* = 5 ms, as the parameters set by the Allen model resulted in very small changes in the Ca^2+^ concentration during APs. Figure 3C and D verifies that the chosen values of *γ* and *τ* yield appropriate changes in [Ca^2+^]_in_ when the neuron is firing [25].

### CaHVA and SK channel activation

The activation curve and time constant of the CaHVA channel is shown in black in Figure 1A and B, respectively. As *τ*_*m*_ *≈* 6 ms at the membrane potential where *m*_*∞*_ = 0.5, it will take the channel some time to open.

**Fig. 1.**
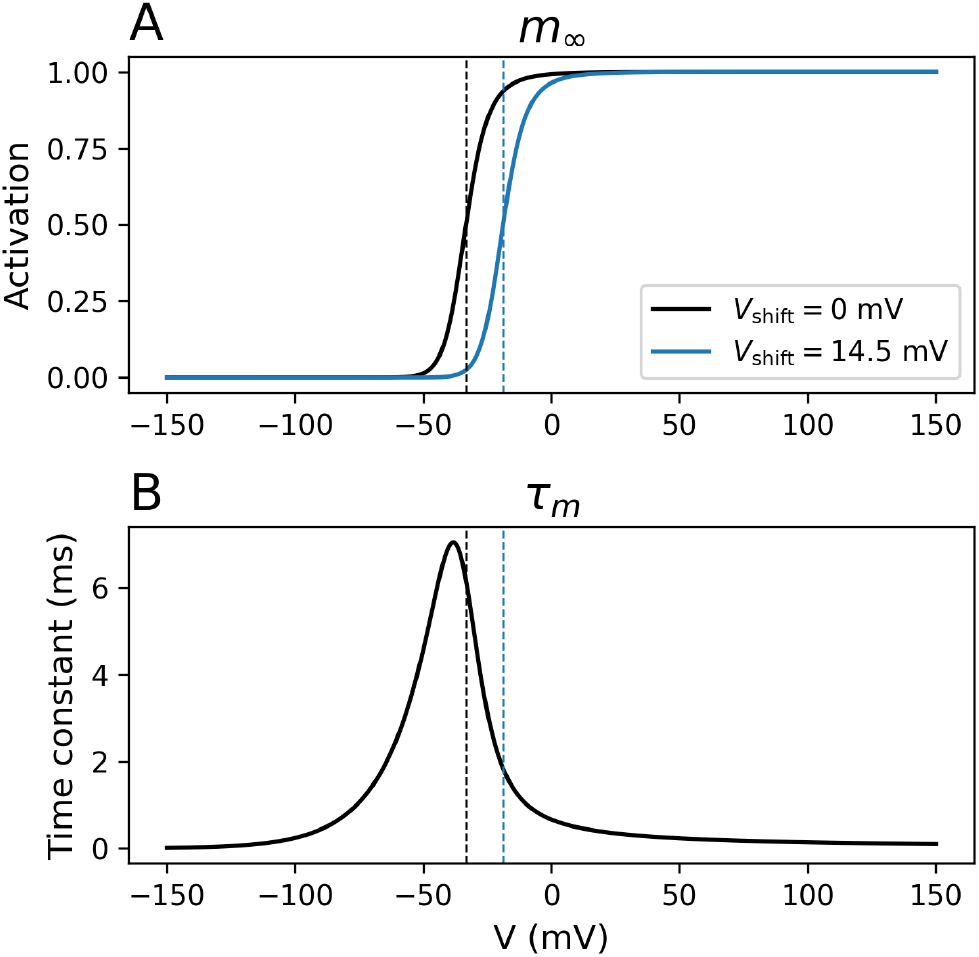
Variables of the CaHVA channel as a function of *V*. A) Activation variable *m*_*∞*_ for the unmodified mechanism (black) and when shifted by 14.5 mV according to literature (blue), B) Time constant *τ*_*m*_ of activation. The dashed lines indicate the voltage at which the activation is at half the maximal value for the shifted and unshifted case.

Figure 2 shows the activation curve of the SK channel, which depends on the intracellular Ca^2+^ concentration. The baseline Ca^2+^ concentration is 10^−4^ mM, as indicated by the gray dotted line. The channel is halfway open when the intracellular Ca^2+^ concentration [Ca^2+^]_in_ has increased from its baseline value by a factor of 4.3.

**Fig. 2.**
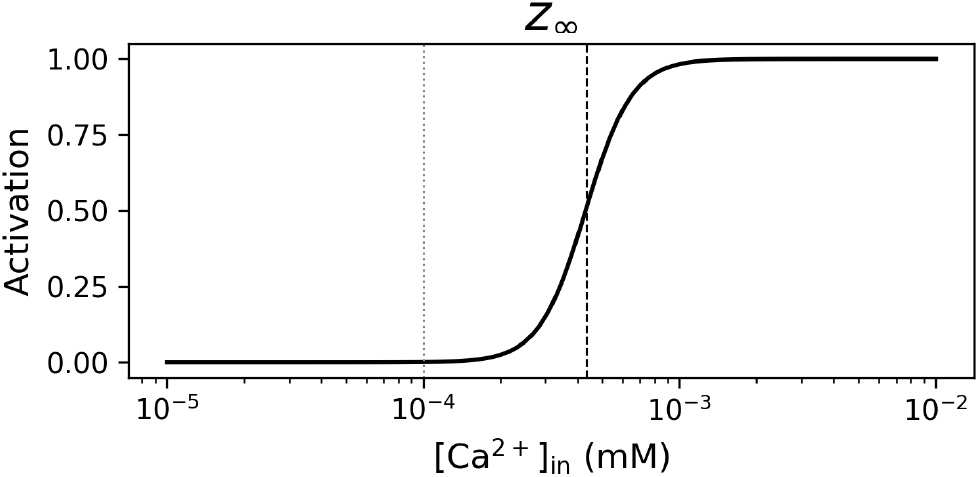
Activation variable *z*_*∞*_ of the SK channel as a function of intracellular Ca^2+^ concentration [Ca^2+^]in. The gray dotted line indicates the baseline value of [Ca^2+^]_in_ in the cell model, while the black dashed line indicates the concentration at which the channel is halfway open.

**Fig. 3.**
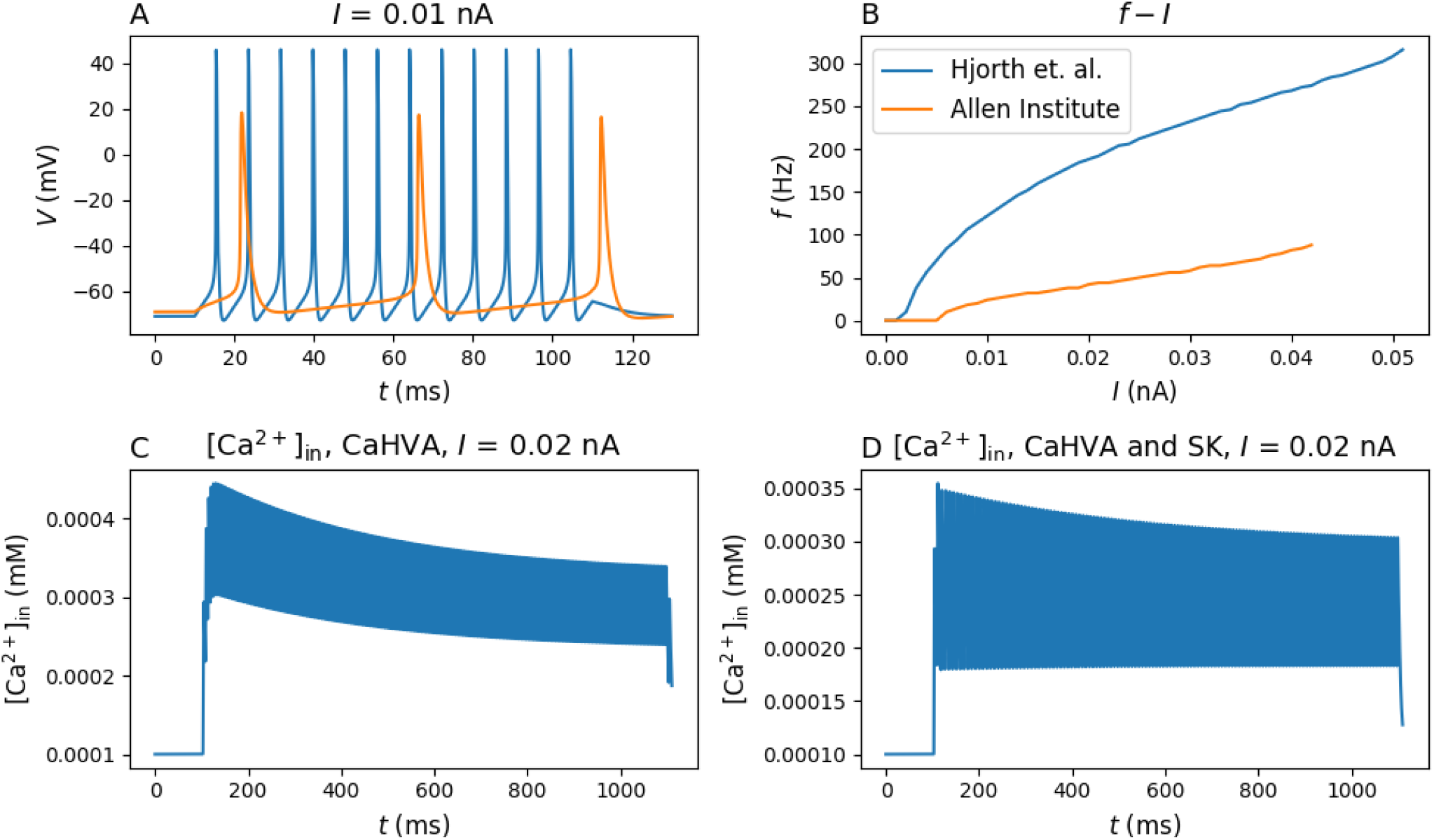
Firing behavior of minimal models of PV cells. A) Voltage traces for input current *I* = 0.01 nA. B) *f* −*I* curve for the minimal models. C) [Ca^2+^]_in_with the minimal Hjorth model 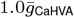 and 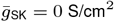 = 0 S/cm, *I* = 0.2 nA. D) [Ca^2+^]_in_ with minimal Hjorth model, 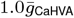and 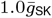*I* = 0.2 nA. The minimal models consist of the following channels: Hjorth et. al. - naf, kaf and leak, Allen Institute - NaV, Kv2like and leak.

### Shifts of the CaHVA channel activation curve

In order to assess the impact of PNN degradation on neuronal firing properties, a shift was introduced in the activation variable of the CaHVA mechanism. The shift was made by performing the substitution *V→ V* −*V*_shift_ in Equation 7 and 8. For completeness, a range of shifts were assessed, from large negative shifts to large positive shifts. The shift observed by Vigetti et al. [32] is shown in blue in Figure 1.

### Additional models

Besides the chosen base model with Na^+^- and K^+^ channels naf and kaf from Hjorth et al. [18], another base model candidate with ion channels NaV and Kv2like from the Allen Institute for Brain Science’s PV neuron models [12] was tested on a one-compartment morphology in our original search for our base model. This candidate base model yielded broader APs and lower firing frequencies than the chosen model with naf and kaf, as shown in Figure 3A and B. As PV cells are known to display a narrow AP duration [2], the base model with naf, kaf and leak was therefore chosen to form the base model that yielded spiking characteristics similar to PV cells.

In addition to ion channels CaHVA and SK taken from the Allen Institute for Brain Science [12], other voltage-gated Ca^2+^ channels and Ca^2+^-activated K^+^ channels were tested to check whether the findings of our main model could be generalized. The results are shown in Supplementary Note A.

### Implementation details

The membrane potential of a one-compartment neuron model was simulated using the NEURON simulation software [4] with LFPy [13] as a wrapper, and analyzed through custom scripts.

The systems were equilibrated for 600 ms before the recordings started. A time step of 0.0078125 ms was used. Input currents *I* of constant amplitude were applied for 1000 ms. Due to varying degrees of spike adaptation, only data from the last 500 ms of the stimulus were analyzed in order to yield readily interpretable results.

## Results

### 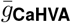 has a limited direct effect on *f*

As seen from Figure 4A, *f* increases with increasing 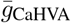 when 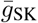 is set to zero. However, this increase is rather small, and firing ceases when 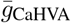 is sufficiently large. Evidently, the CaHVA conductance does not increase *f* on its own.

**Fig. 4.**
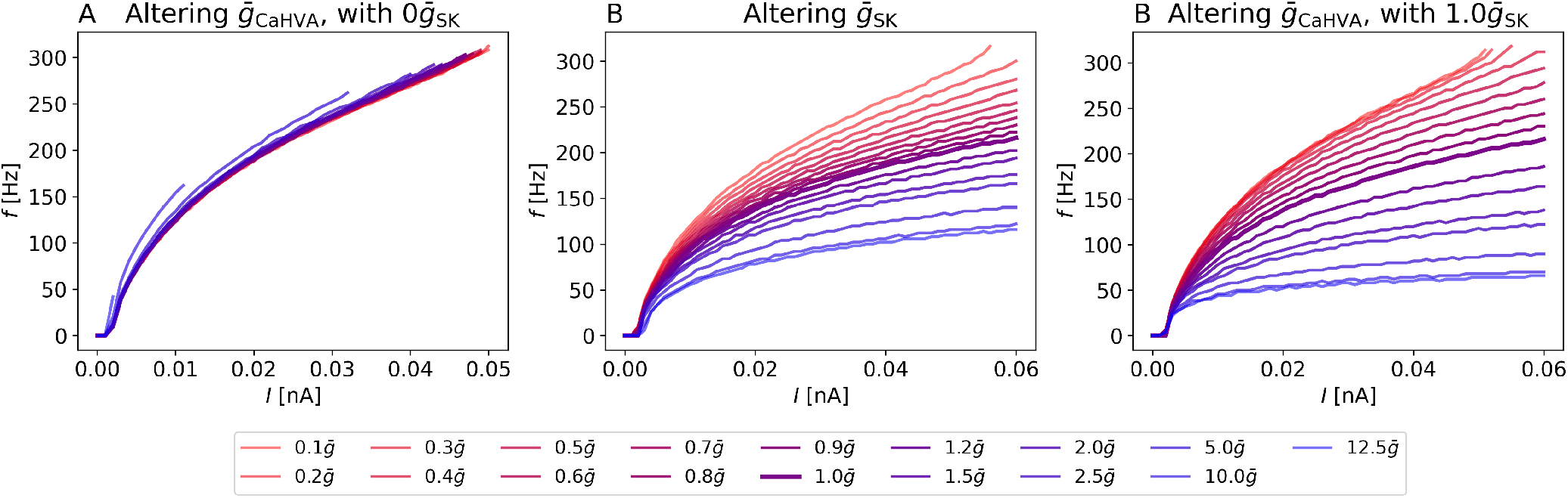
*f* − *I* curves when altering A) 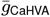 with the SK conductance set to zero, B) 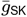 with default 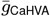, C) 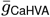with the SK conductance set to default 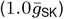 Bold lines represent curves of the default parameters for each of the models.

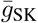, on the other hand, displays a larger effect on *f*, as shown in Figure 4B where *f* is shown to decrease markedly with increasing 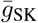. However, it requires that CaHVA is present.

### 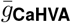 affects *f* mainly via activation of SK

Figure 4C reveals that when 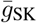 is non-zero, increasing 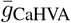 has the opposite effect on firing than when 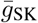 is zero. In this case, *f* decreases drastically with increasing 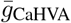. This is because an increase in 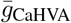 increases the intracellular Ca^2+^ concentration, which leads to a stronger activation of the SK channel. In turn, this leads to a membrane hyperpolarization after the action potential due to the positive K^+^ current *I*_SK_. The lowered membrane potential makes it harder for the neuron to fire, which is reflected in the reduced firing rates.

However, 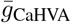 is implied to decrease upon PNN removal [20], which should lead to an increase in *f* according to Figure 4C. Therefore, the reduction in *f* observed experimentally [1, 28, 22, 5] is not likely to be caused by a change in 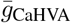.

For completeness, we tested other CaHVA channels and Ca^2+^-activated K^+^ channels SK or BK from the literature on our base model. The conductances were varied in the same manner as above. The results were qualitatively similar to the main model in that the primary effect of increasing 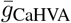 was indirect, i.e. through activation of SK. The results are shown in Figure 10 in Supplementary Note A.

### Shifting activation curves

#### Shifts in CaHVA activation curves have little direct effect on f

Vigetti et al. [32] found that the presence of CS in solution induced a shift in the CaHVA activation curve rather than 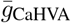. We therefore tested how various shifts in CaHVA affected *f*. Figure 5 shows *f* vs *I* for values of *V*_shift_ ranging from large and negative to large and positive, shifting the activation curves to lower and higher voltages, respectively. When 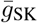 is set to zero, as seen in Figure 5A-D, negative shifts lead to an increase in *f*, consistent with CaHVA channels being open for a larger range of voltages and consequently having a larger effect. Contrarily, positive shifts lead to a reduction in *f*, although this reduction is limited. At sufficiently large positive shifts, the CaHVA channel will activate at voltages outside the dynamic range of the membrane potential, so that a further increase in *V*_shift_ has no additional effect. PNN degradation is associated with positive shifts [32]. Similarly to 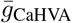 an increase in *V*_shift_ has little direct effect on *f*, as seen from Figure 5A-D.

**Fig. 5.**
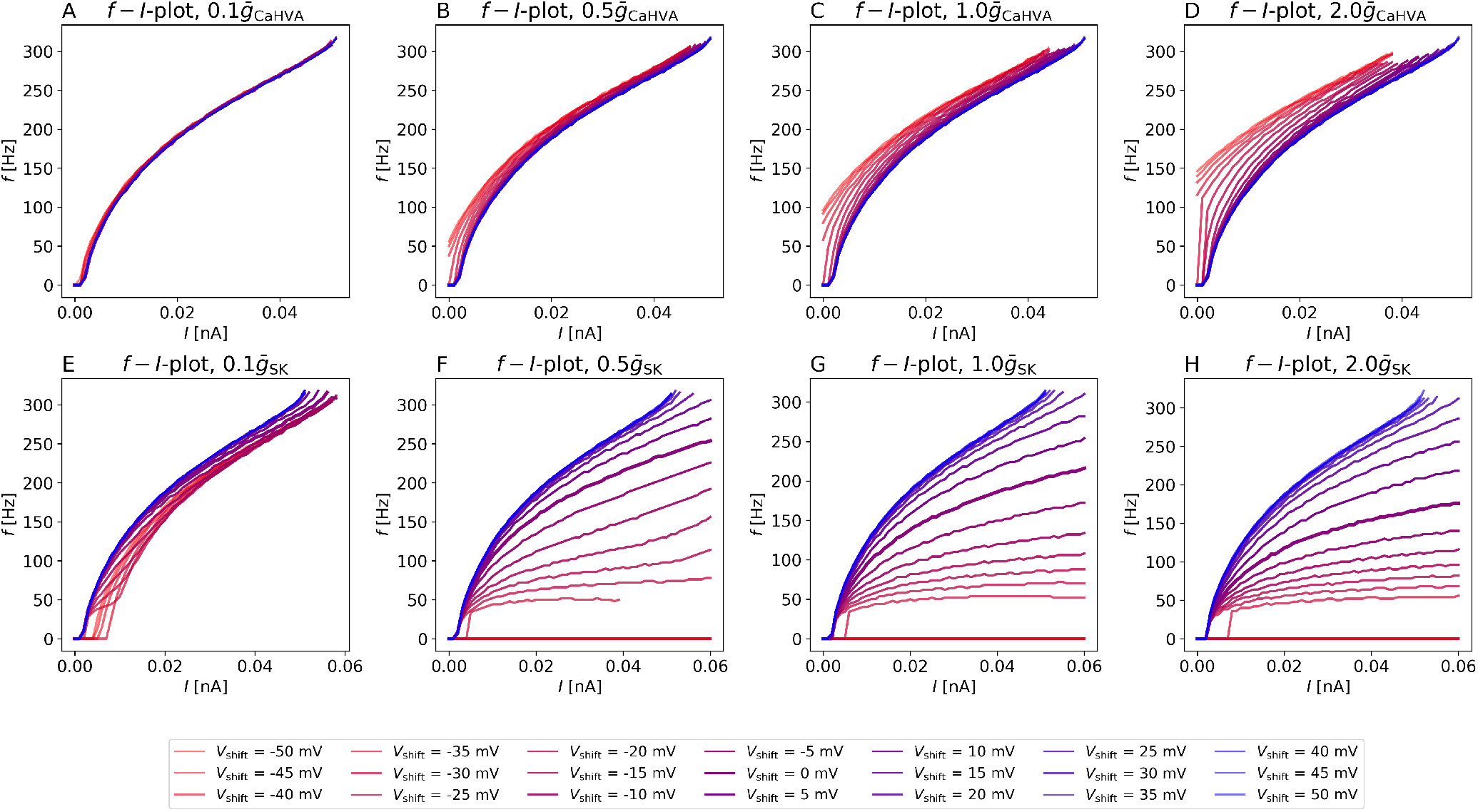
*f I* curves for different shifts *V*_shift_ of the CaHVA activation curve. A) 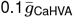, B) 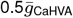, C) 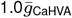, D) 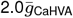E) 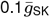, F) 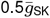 G) 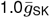, D) 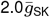. In A)-D), 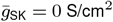.

As PNN degradation is associated with positive shifts [32], these results suggest that reduced firing in the absence of the nets (as observed by e.g. Tewari et al. [28]) could at least partially be due to the effect of PNNs on CaHVA channel activation, although this effect has a limited impact on *f*.

As seen from Figure 5A-D the effect of the shifts varies depending on the value of 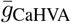. We denote an increase in the maximal conductance by a factor of *a* by 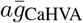 for brevity, meaning that 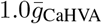 is the default value. For 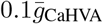, the effect of *V*_shift_ on *f* is rather small, with the effect of positive shifts being negligible. For 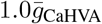, the effect of the shift becomes more prominent. Negative shifts have a larger effect. However, the largest negative shifts yield spontaneous firing (i.e. shifts towards lower voltages, opposite of PNN removal). The spontaneous firing is a side effect of the CaHVA channel being open at the resting membrane potential when *V*_shift_ is lowered sufficiently.

### Shifts in CaHVA activation affect f via activation of SK

While a shift in CaHVA has little effect on *f* on its own, its influence on *f* through activation of the SK current is large. Figure 5E-H shows *f* vs *I* for the selected values of *V*_shift_ when the small-conductance Ca^2+^-activated K^+^ channel SK is present. Comparing with Figure 5A-D, *f* responds greatly to shifts in CaHVA channel activation. No spontaneous firing occurs for any combination of *V*_shift_ and 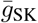. There is a slight change in the onset of firing for small 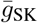as seen from Figure 5E. For larger values of 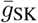, the neuron eventually stops firing when a negative *V*_shift_ is imposed.

A negative shift in CaHVA channel activation leads to a decrease in *f* for all 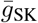 while a positive shift leads to an increase in *f* for all 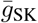. Since PNN-removal is associated with a positive shift, these results suggest that the shift in activation could not contribute to the reduced firing observed by in PV cells upon PNN removal [1, 22, 5].

Due to the interplay between CaHVA and SK currents, *V*_shift_ can alter *f* either directly or indirectly. Judging by Figure 5, the largest impact of *V*_shift_ on firing is indirect through its effect on *I*_SK_.

### A mechanistic understanding of the *f*-regulation

Figure 6A shows the trace over one action potential when varying 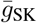 with 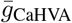 set to default. There are minute changes to the AP shape when 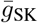 is varied, but the most pronounced effect is that *V* lingers at a slightly lower value after the AP, as seen from Figure 6C. A larger rise in the membrane potential is therefore required in order to reach the action potential threshold when 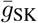 is increased, which means that *f* should decrease with increasing 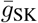. These observations are in agreement with the *f* −*I* curves of Figure 4B, and SK currents being classified as afterhyperpolarizing [9].

**Fig. 6.**
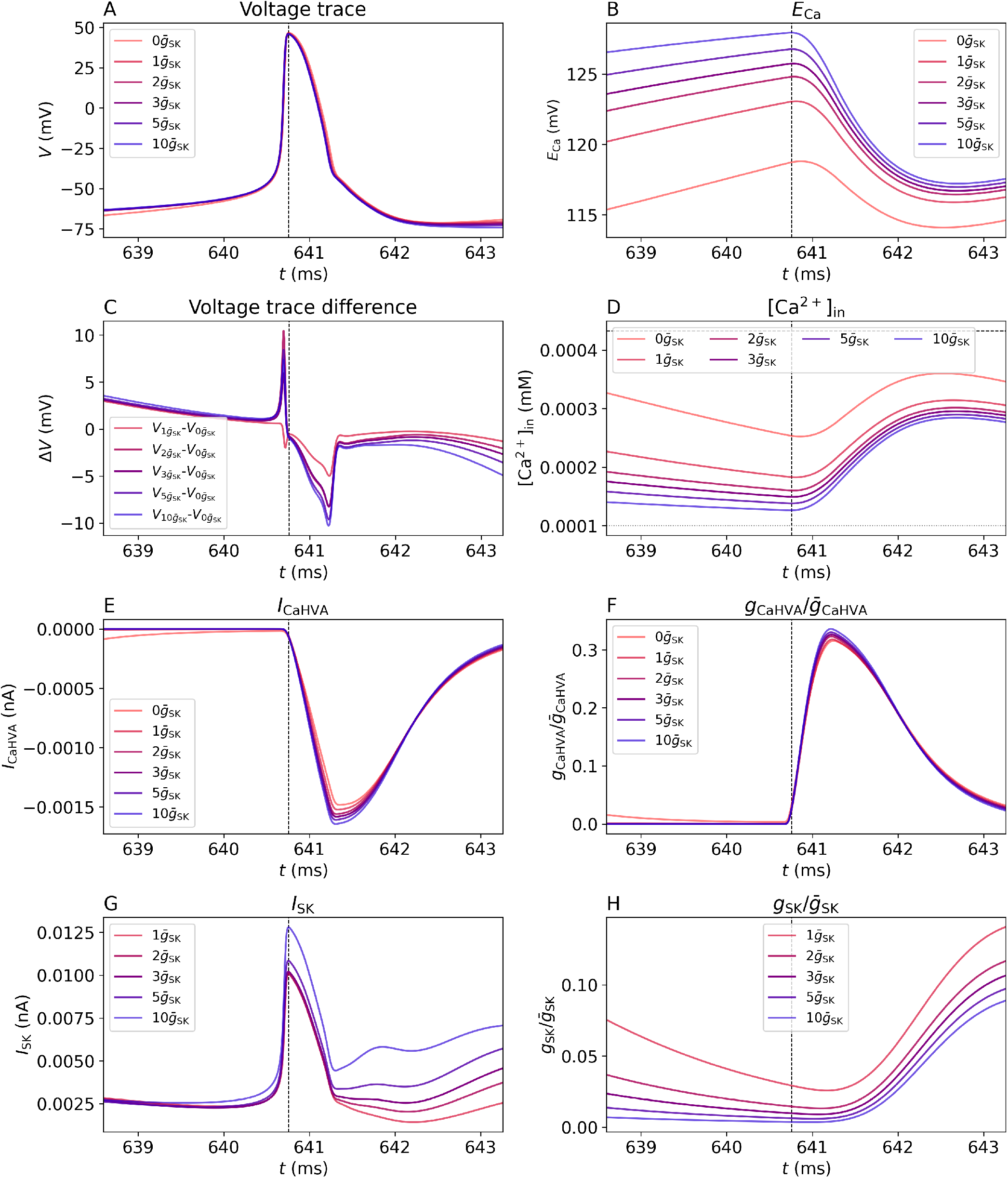
A) Voltage trace over one AP when varying 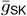. B) *E*_Ca_. C) Difference between voltage traces with and without the SK channnel. D) [Ca^2+^]_in_. The dotted line is the initial concentration and the dashed line is the concentration at which the channel is halfway open. E) *I*_CaHVA_. F) 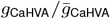. G) *I*_SK_. H) 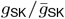. The peaks have been aligned to allow for easy comparison between models with different values of 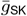. The time of the action potential peak is indicated by the vertical dashed line.

*I*_SK_ is determined by three things: the conductance *g*_SK_, the intracellular Ca^2+^ concentration [Ca^2+^]_in_ and the driving force *V* −*E*_K_. During different phases of the firing, different variables dictate the value of *I*_SK_. For the duration of the AP, *g*_SK_ and [Ca^2+^]_in_ are nearing minima (see Figure 6D and 6H), with *g*_SK_ being small, but non-zero. The increase in *I*_SK_ seen in Figure 6G must therefore be primarily due to the increase in driving force as *V* changes in Figure 6A. After the AP, the increases in *g*_SK_ and [Ca^2+^]_in_ are responsible for the rise in *I*_SK_ as *V* and hence *V*− *E*_K_ decreases in this interval.

The Ca^2+^ current is initiated just before the peak of the action potential, and becomes large and negative in the following few milliseconds, as seen from Figure 6E. Correspondlingly, the relative Ca^2+^ conductance 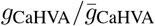, shown in Figure 6F, starts increasing right before the peak of the action potential, rises rapidly and decays slowly. Evidently, the CaHVA channel mainly opens during the later phase of the AP, consistent with CaHVA being a high-voltage-activated channel and having a large time constant of activation *τ*_*m*_ (see Figure 1B). The channel then stays open for a while after the action potential, consistent with the large value of *τ*_*m*_. The CaHVA channel will thereby mainly affect the value of the membrane potential after the peak, as this is when current flows through the channel. If the membrane potential stabilizes at a higher value, it will be closer to the firing threshold, and less current will be needed to intiate a new AP, while more current will be needed if it stabilizes at a lower value. Therefore, the CaHVA channel can affect how fast the neuron is able to fire.

When the shift in CaHVA channel activation is introduced and the SK conductance is non-zero, the membrane potential increases with increasing *V*_shift_ after the AP, as seen from Figure 7A. This is reflected by the membrane potential difference shown in 7C, and should make it slightly easier for the neuron to fire as *V* is closer to the firing threshold. A positive *V*_shift_ should therefore increase *f*, as we saw in Figure 5.

**Fig. 7.**
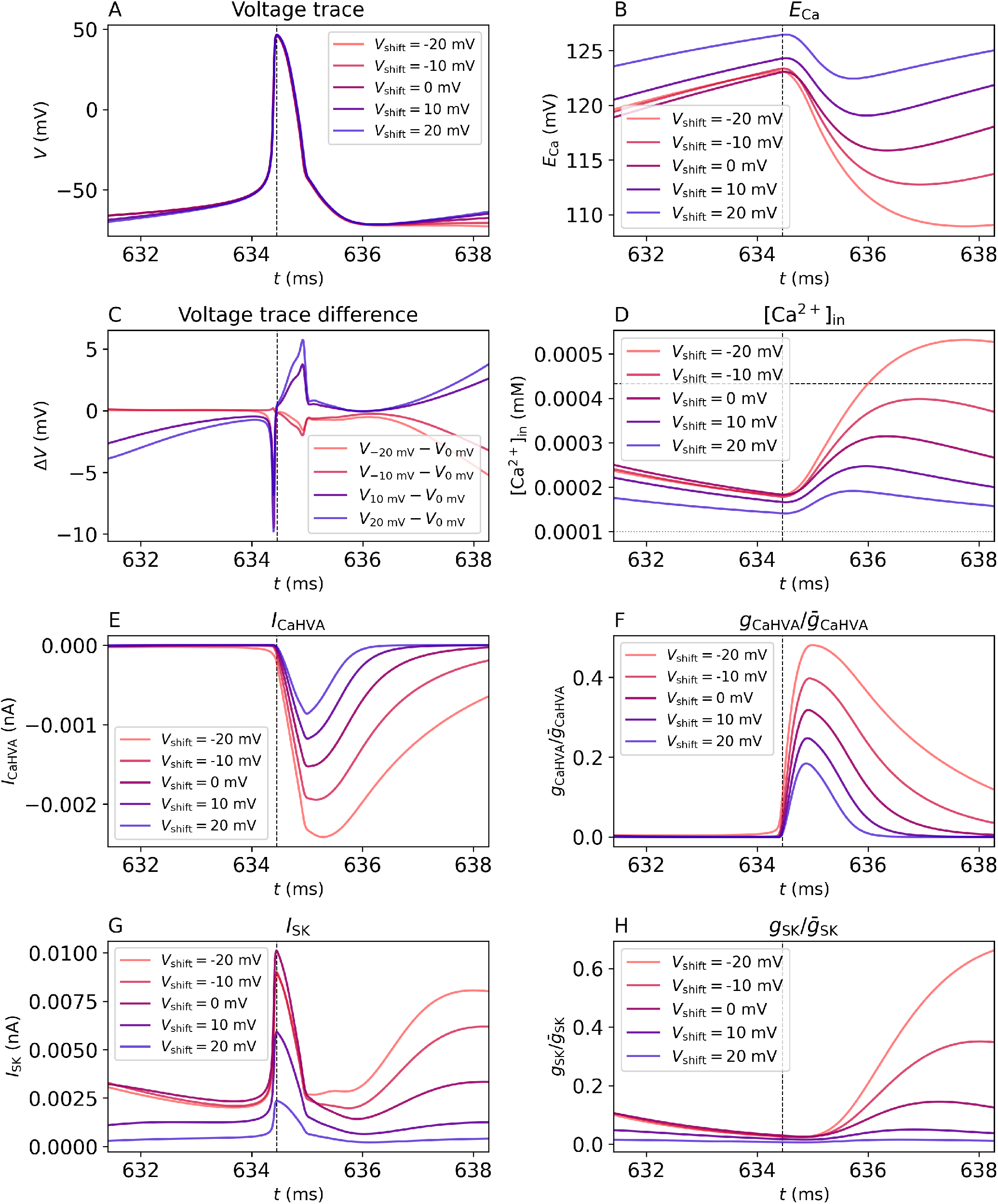
A) Voltage trace over one AP for various *V*_shift_ when CaHVA and SK channels are present. B) *E*_Ca_. C) Difference between voltage traces with and without the shift. D) [Ca^2+^]_in_. The dotted line is the initial concentration and the dashed line is the concentration at which the channel is halfway open. E) *I*_CaHVA_. F) 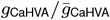. G) *I*_SK_. H) 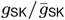. The peaks have been aligned to allow for easy comparison between models with different values of *V*_shift_. The time of the action potential peak is indicated by the vertical dashed line.

The primary effect of *V*_shift_ is to determine the membrane potential at which the CaHVA channel opens. As can be seen from studying Figure 7F thoroughly, *g*_CaHVA_ starts to rise a little bit sooner for *V*_shift_ = −20 mV than for default. This is consistent with the shifted channel activating at a lower membrane potential. For *V*_shift_ = − 20 mV, the channel starts to open before the abrupt rise in *V* during depolarization. For the other values of *V*_shift_, the channel opens during the depolarization phase, so that the onset of the rise in 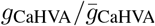 is not sensitive to *V*_shift_.

The size of 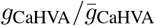 does however depend on *V*_shift_. As can be seen from Figure 1 and Eq. 4, *g*_CaHVA_ can reach higher values when the CaHVA channel activation is shifted towards lower voltages. As a consequence, both *g*_CaHVA_ and *I*_CaHVA_ increase in absolute magnitude with decreasing *V*_shift_, as seen in Figure 7E and F.

A consequence of the increased *I*_CaHVA_ with decreasing *V*_shift_ is the increase in [Ca^2+^]_in_ with decreasing *V*_shift_ displayed in Figure 7D. This, in turn, affects the Ca^2+^-activated SK channel. Figure 7G and H reveals that *I*_SK_ and 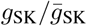 are consistently larger for negative *V*_shift_, as more intracellular Ca^2+^ is present. This holds even after the AP, lowering *V* in Figure 7A compared to default for *V*_shift_ *<* 0 mV. *V* therefore has to increase more in order to reach the firing threshold when *V*_shift_ *<* 0 mV, which should lower *f*. This is exactly what we have observed in Figure 5.

### Changes in parameter values consistent with PNN degradation alter *f* in different ways in different model implementations

Vigetti et al. [32] found that removal of PNN component chondroitin from solution caused a 14.5 mV shift in the activation of voltage-gated Ca^2+^ channels. This shift should lead to a lower [Ca^2+^]_in_ as the Ca^2+^ channels activate at higher voltages. In turn, this should make it harder for the SK channel to activate, causing less *I*_SK_ and a higher *f*.

Dembitskaya et al. [6] found a 3.337 factor increase in 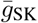 upon PNN removal with chABC. This should increase *I*_SK_ and lower *f* as a consequence. As the observed changes in CaHVA and SK associated with PNN removal affect *f* in opposite ways, it is not obvious how *f* will be altered by PNN breakdown. The exact strength and form of the channels used in the computational model could also affect which mechanism dominates the firing rate.

Using the 14.5 mV shift in CaHVA activation and the 3.337 factor increase in 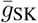 our main model displayed an increase in *f*, as seen in Figure 8. This is opposite to what is reported in the literature [1, 28, 22, 5], meaning that according to the main model the observed PNN-induced changes in the CaHVA and SK channels cannot concomitantly contribute to the reduced firing found in these experiments.

**Fig. 8.**
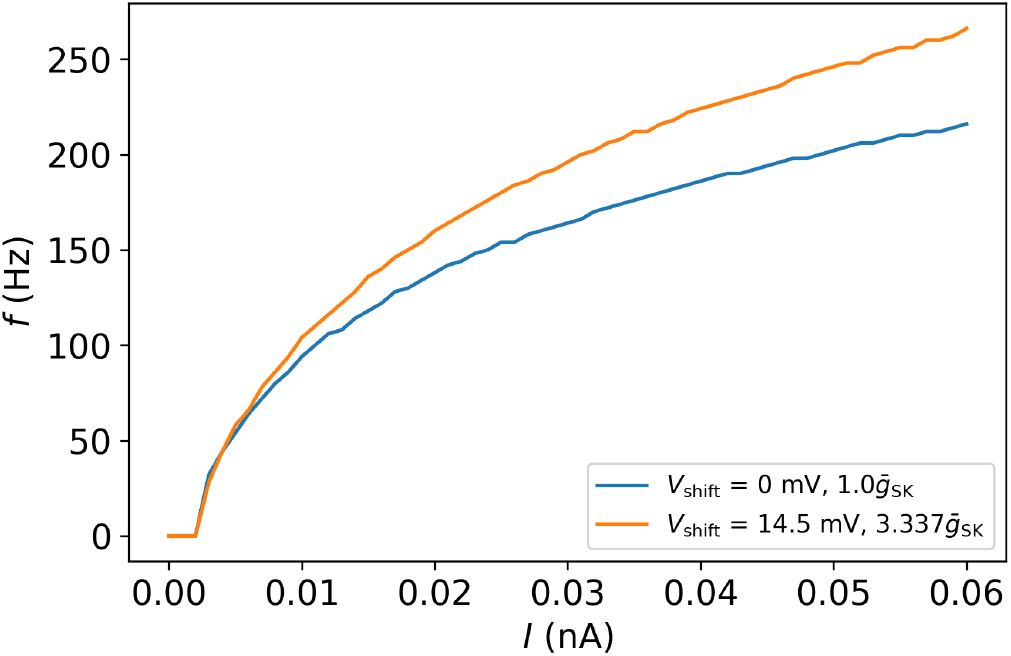
*F*− *I* curves for the default model (*V*_shift_ = 0 mV, 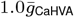and and 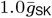) and the model that is altered to be consistent with PNN degradation (*V*_shift_ = 14.5 mV, 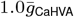 and 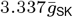). 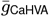 and 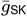 are the default conductances as defined in the Methods section.

Other choices of CaHVA and Ca^2+^-activated K^+^ channels displayed different results when these realistic alterations were performed. The different models displayed different qualitative responses to the parameter value changes, ranging from a reduction in *f*, to no change in *f*, and to an increase in *f*. The results are shown in Figure 9. For most of the models, the changes in the *f*− *I* curves are quite small. The exception to this are the models with BK from Hjorth et al. [18], which display a large shift when the parameter values are altered according to the literature. Quite a few of the models display a reduction in *f*, which is consistent with literature on the effect of PNN breakdown on *f* [1, 28, 22, 5]. This means that the changes to CaHVA and SK channels could help explain the observed reduction in *f*, but our results remain somewhat inconclusive.

**Fig. 9.**
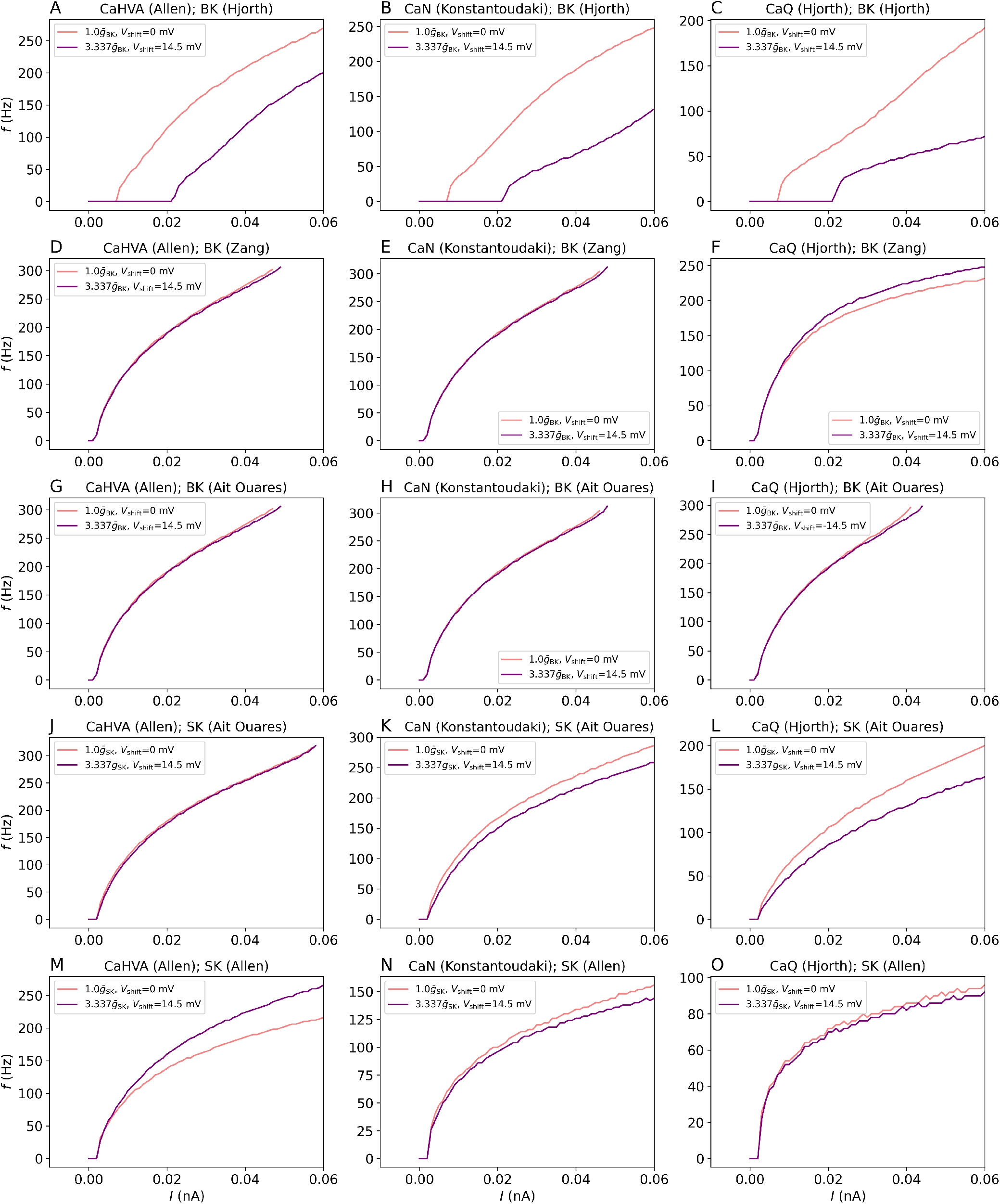
*f*− *I* curves for the default model (*V*_shift_ = 0 mV, 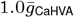and and 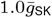) and the model that is altered to be consistent with PNN degradation (*V*_shift_ = 14.5 mV, 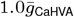 and 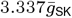). A) CaHVA (Allen) and BK (Hjorth), B) CaN (Konstantoudaki) and BK (Hjorth), C) CaQ (Hjorth) and BK (Hjorth), D) CaHVA (Allen) and BK (Zang), *E)*CaN (Konstantoudaki) and BK (Zang), F) CaQ (Hjorth) and BK (Zang), G) CaHVA (Allen) and BK (Ait Ouares), H) CaN (Konstantoudaki) and BK (Ait Ouares), I) CaQ (Hjorth) and BK (Ait Ouares), J) CaHVA (Allen) and SK (Ait Ouares), K) CaN (Konstantoudaki) and SK (Ait Ouares), L) CaQ (Hjorth) and SK (Ait Ouares), M) CaHVA (Allen) and SK (Allen), N) CaN (Konstantoudaki) and SK (Allen), O) CaQ (Hjorth) and SK (Allen). The cell models consist of the channels mentioned as well as channels naf and kaf from Hjorth et al. and a standard leak channel. The default parameters are listed in Table 1.

**Fig. 10.**
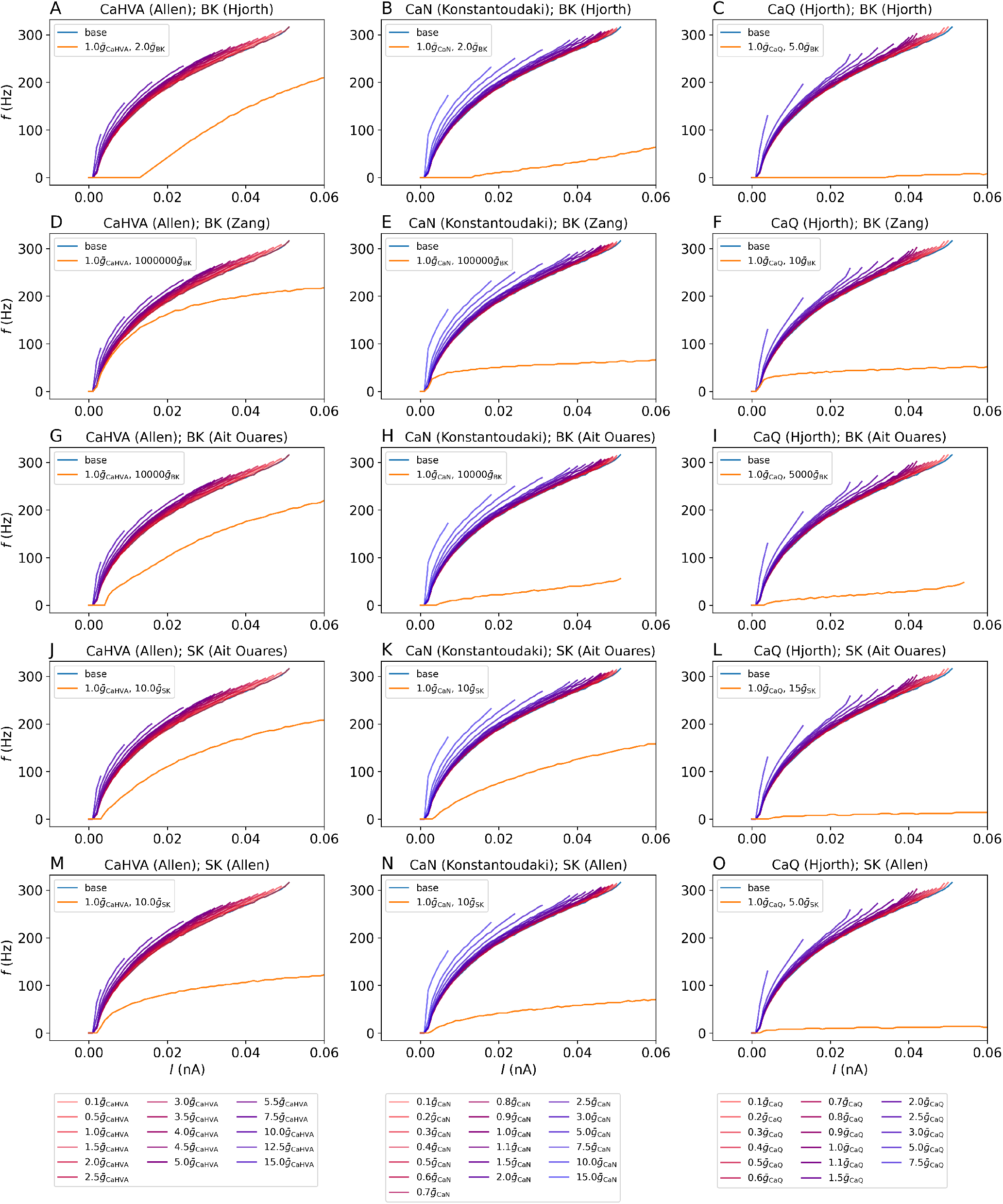
*f* −*I* curves for different combinations of ion channel models over a range of Ca^2+^ conductances without SK or BK currents present, as well as for one value of 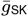or 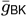. A) CaHVA (Allen) and BK (Hjorth), B) CaN (Konstantoudaki) and BK (Hjorth), C) CaQ (Hjorth) and BK (Hjorth), D) CaHVA (Allen) and BK (Zang), E) CaN (Konstantoudaki) and BK (Zang), F) CaQ (Hjorth) and BK (Zang), G) CaHVA (Allen) and BK (Ait Ouares), H) CaN (Konstantoudaki) and BK (Ait Ouares), I) CaQ (Hjorth) and BK (Ait Ouares), J) CaHVA (Allen) and SK (Ait Ouares), K) CaN (Konstantoudaki) and SK (Ait Ouares), L) CaQ (Hjorth) and SK (Ait Ouares), M) CaHVA (Allen) and SK (Allen), N) CaN (Konstantoudaki) and SK (Allen), O) CaQ (Hjorth) and SK (Allen). The cell models consist of the channels mentioned as well as channels naf and kaf from Hjorth et al. and a standard leak channel. The default parameters are listed in Table 1.

Note that some of the models in Figure 9 include the BK channel instead of the SK channel, which complicates the interpretation of the parameter value change as the observations were made for the SK conductance in the literature [6].

These models are still included for completeness.

Comparing Figure 10 to Figure 9, it becomes clear that the models that required a large 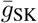 in order to display a reduced firing (panels D, E, G, H and I) all show little to no change in firing upon a realistic alteration of 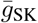 and *V*_shift_. The remaining models, where *f* was more sensitive to 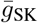 for the most part displayed a larger difference in *f*. This observation supports the notion that the SK current is mainly responsible for the change in *f*.

Referring to Figure 9, the CaHVA channel used by the Allen Institute for Brain Science either causes an increase in *f* when 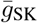 and *V*_shift_ is increased according to the literature, no alteration, or a decrease and a large shift when BK from [18] is present. Models including the CaN channel for inhibitory neurons from [21] always display either no change in *f* or a decrease in *f*. The CaQ channel from [18] mostly displays a reduction in *f*, but also one case of an increase and one case with no change in *f*.

## Discussion and conclusion

We studied the interplay between SK and CaHVA channels in computational models of PV neurons. Altering the CaHVA current when 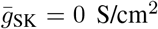, be it by increasing the conductance or shifting the activation curve to lower values, leads to a minor increase in *f*. When 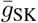 is non-zero, CaHVA currents will trigger *I*_SK_ due to rises in the intracelluar Ca^2+^ concentration. An increase in 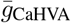 will increase [Ca^2+^]_in_ more strongly, leading to a larger *I*_SK_. As *I*_SK_ is a repolarizing current, *V* will linger at lower values after the AP, and a larger increase in *V* will be needed to reach the firing threshold. *f* therefore decreases with increasing 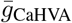 through activation of *I*_SK_. This indirect decrease in *f* is more pronounced than the direct decrease in *f* that was observed when 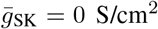. We consider this to be a general finding, as it was found to hold for all models tested, as shown in Supplementary Note A.

When 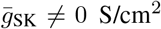, positive shifts in CaHVA channel activation leads to a decrease in *I*_SK_, causing an increase in *f*. As an increase in 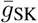was also shown to lead to an increase in *I*_SK_ and *f*, we conclude that the impact on *f* from changing 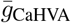 *V*_shift_ or 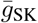 ultimately depends on their effect on *I*_SK_.

We altered *V*_shift_ and 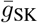 according to literature on neurons with and without PNNs, and found that the response of *f* to PNN breakdown depends on the choice of model. In some cases, *f* was reduced when these parameter changes where made.

Note that CaHVA channels come in several types that differ in behavior [33]. The high-voltage-activated Ca^2+^ channels are divided into L-, N-, P-, Q-, and R-type [8]. All of these have been found on PV neurons [33]. The type of the CaHVA channel primarily used in this work is unspecified, while channels like CaN borrowed from Konstantoudaki et al. [21] and CaQ borrowed from Hjorth et al. [18] are more specific.

The interplay between voltage-activated Ca^2+^ channels and Ca^2+^-activated K^+^ channels have previously been studied in computational models of cells in the heart [19], pituitary cells [14] and in the brain [11, 15, 23, 26].

For simplicity, we limited our study to one-compartment models. Different results could have been obtained by using a complex morphology. However, then we would have needed detailed information on e.g. distribution of SK channels, which is normally not available. Since the aim of this work was to show the interplay between high-voltage-activated Ca^2+^ channels and Ca^2+^-activated K^+^ channels, emphasis was put on testing out different channels instead of exploring morphologies. As qualitative results are more readily interpretable in a simplified model, an investigation of CaHVA and SK or BK channels in complex morphologies has been left to future works.

## Code availability

The code used to produce and analyze the results is available on Github: https://github.com/KineOdegardHanssen/CaHVA_SK_PNN_code.

## Supplementary Note A: Generalization to other channels

In order to test if our findings are general, other relevant ion channels were tested on the base model introduced in the Methods section. The models were taken from Hjorth et al. [18], Konstantoudaki et al. [21], Zang et al. [35], Ait Ouares et al. [26] and the Allen Institute for Brain Science [12].

As the ion channels were taken from models with advanced morphologies and put on a one-compartment model, some of the conductances were changed from their values in the original works. 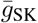 and 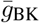 were set to the value found in the Allen model. The value of 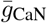 was kept to its original value from Konstantoudaki et al. [21], while 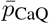 was adjusted to yield an *f* −*I* curve similar, but not identical to the base model. A list of the default conductances of this paper is provided in Table 1. Similar to the *f*−*I* curve of Figure 4A, if no SK or BK current is present, all neuron models stop firing when the Ca^2+^ conductance is sufficiently increased, as seen by Figure 10. Furthermore, Figure 10 shows that for all models, the firing rate decreases significantly for a sufficiently large SK or BK conductance. Although the magnitude of this reduction and the required SK or BK conductance is model-dependent, the finding is in qualitative agreement with Figure 4B.

**Table 1.** Default conductances used in Figure 10. Note that some conductances have been changed a bit due to the change in morphology.

| $\bar{g}_{\text{CaHVA}}$ (S/cm<sup>2</sup>) | $\bar{g}_{\text{CaN}}$ (S/cm<sup>2</sup>) | $\bar{p}_{\text{CaQ}}$ (cm/s) | $\bar{g}_{\text{SK}}$ (S/cm<sup>2</sup>) | $\bar{g}_{\text{BK}}$ (S/cm<sup>2</sup>) |
| --- | --- | --- | --- | --- |
| $2.99 \cdot 10^{-5}$ | 0.0003 | $6.2 \cdot 10^{-5}$ | 0.0028 | 0.0028 |

## Notes

https://github.com/KineOdegardHanssen/CaHVA_SK_PNN_code

### Competing Interest Statement

The authors have declared no competing interest.

